# *GABRD* promotes the progression of breast cancer through CDK1-dependent cell cycle regulation

**DOI:** 10.1101/2023.10.10.561812

**Authors:** Qingyao Shang, Fei Ren, Kexin Feng, Chenxuan Yang, Shuangtao Zhao, Jiaxiang Liu, Xiyu Kang, Jiaxian Yue, Ruixuan Zhang, Xiangzhi Meng, Xiang Wang, Xin Wang

## Abstract

**Purpose:** Y-aminobutyric acid (GABA) is an important inhibitory amino acid neurotransmitter that exerts its biological function by binding to GABA receptors, which not only play an important role in neuromodulation, but also involved in regulating the development of tumors. Gamma-aminobutyric acid type A receptor subunit delta (*GABRD*) encodes the δ subunit of GABA_A_ receptor, its impact on breast cancer has not been clearly studied. This study is aiming to reveal the relationship between *GABRD* and breast cancer development.

**Methods:** We performed a tissue microarray to quantify *GABRD* expression levels in tumor tissue and paracarcinoma tissue. The regulation of *GABRD* in the proliferation, migration, and apoptosis of breast cancer was examined by a loss-of-function study. A GeneChip microarray was used to probe GABRD for potential downstream molecules. The interaction between GABRD and CDK1 was verified by a set of functional tests and rescue experiments as well as coimmunoprecipitation.

**Results:** *GABRD* was expressed at significantly higher levels in tumor tissues and was associated with advanced tumor progression. Silencing *GABRD* resulted in a significant decrease in proliferation and migration and an increase in apoptosis of breast cancer. *GABRD* regulated the cell cycle by directly interacting with CDK1, which was identified as an important downstream target.

**Conclusion:** *GABRD* is the breast cancer-related gene and highlights the importance of the GABRD–CDK1 axis in regulating breast cancer proliferation, which provides potential for the development of novel therapeutics.

## Introduction

Breast cancer is the most common female malignancy and the leading cause of mortality worldwide, with approximately 2.26 million new cases in 2020, making breast cancer the world’s most prevalent cancer(1). Currently, breast cancer treatment mainly consists of the combination of surgery, chemotherapy, radiotherapy and hormonal therapy. The precise treatment of tumors by molecular-targeted therapy has recently appeared to be an important method, providing additional therapeutic options for several molecular subtypes of breast cancer and improving treatment outcomes. However, breast cancer treatment is limited by the lack of more effective targets and drugs. Therefore, identifying novel and effective biomarkers may improve breast cancer prognoses and provide new therapeutic strategies.

As the predominant and most prevalent inhibitory neurotransmitter in the central nervous system, γ-aminobutyric acid (GABA) is involved in GABAergic signaling to mediate the synaptic inhibition of neuronal activity in the mammalian brain(2). GABA transmits signals by binding with its receptors, which mainly include GABA_A_ and GABA_B_ receptors(3). The most closely related receptor, the GABA_A_ receptor, is a pentamer assembled from multiple subunits that form a chloride channel(4). Among them, the δ subunit is encoded by the *GABRD* gene. *GABRD* gene abnormalities have been found to be closely related to spontaneous epilepsy and have attracted the attention of researchers(5). Most previous studies on *GABRD* have focused on neurological diseases, and several studies have recently revealed the possible functional role of *GABRD* in tumors. Studies based on genomics datasets including mRNA expression data from The Cancer Genome Atlas (TCGA) database showed that *GABRD* expression was associated with corticotrophinomas, hepatocellular carcinoma, kidney renal clear cell carcinoma, thyroid carcinoma and other tumors, which indicated that *GABRD* might serve as a potential biomarker to predict patient prognoses(6–8). However, little is known about the involvement of *GABRD* in the tumorigenesis and progression of breast cancer.

Herein, *GABRD* expression levels in breast cancer and normal tissue samples from patients combined with their correlations with clinical–pathological characteristics were analyzed by tissue microarray and immunohistochemistry (IHC). We identified *GABRD* as a key gene in breast cancer tumorigenesis and progression. Then, we knocked down *GABRD* in MDA-MB-231 and BT-549 cell lines through lentivirus-mediated short hairpin RNA (shRNA) silencing. *GABRD* functions in cell proliferation, migration and apoptosis were investigated in the aforementioned cell lines and xenograft mouse models. GeneChip microarray analysis revealed that CDK1 is a critical target protein regulated by *GABRD*, which was verified by gene rescue experiments. Our research aimed to evaluate the role of *GABRD* in breast cancer progression and unveil the potential associated mechanisms.

## Results

### *GABRD* is implicated in breast cancer progression and invasion

Breast cancer expression profile was obtained from the TCGA database and the normal tissue RNA-seq profile was obtained from the TCGA database and GTEx database. By comparing the expression differences of GABA_A_R and GABA_B_R subunits in tumor and normal tissues, it was found that GABRD was highly expressed in breast tumors and the difference was the most significant (Fig. 1A). GABRD expression in breast cancer tissue and normal tissue was then determined by tissue microarrays. Representative images are shown in Fig. 1B. We found that GABRD expression in breast cancer tissue was markedly increased compared with that in paracarcinoma tissue (Fig. 1B, C). A total of 55.6% of tumor tissues showed high *GABRD* expression, while 0% of paracarcinoma tissues showed high *GABRD* expression (Table 1). By combining the IHC analysis results with clinicopathological information, we found that *GABRD* expression possessed a significant correlation with the clinical stage, T-cell infiltration and tumor size (Table 2). Moreover, according to Kaplan–Meier survival analysis, *GABRD* gene expression was significantly related to the overall survival of breast cancer patients, suggesting that high *GABRD* expression predicted a more unfavorable prognosis (Fig. 1D).

**Figure 1.**
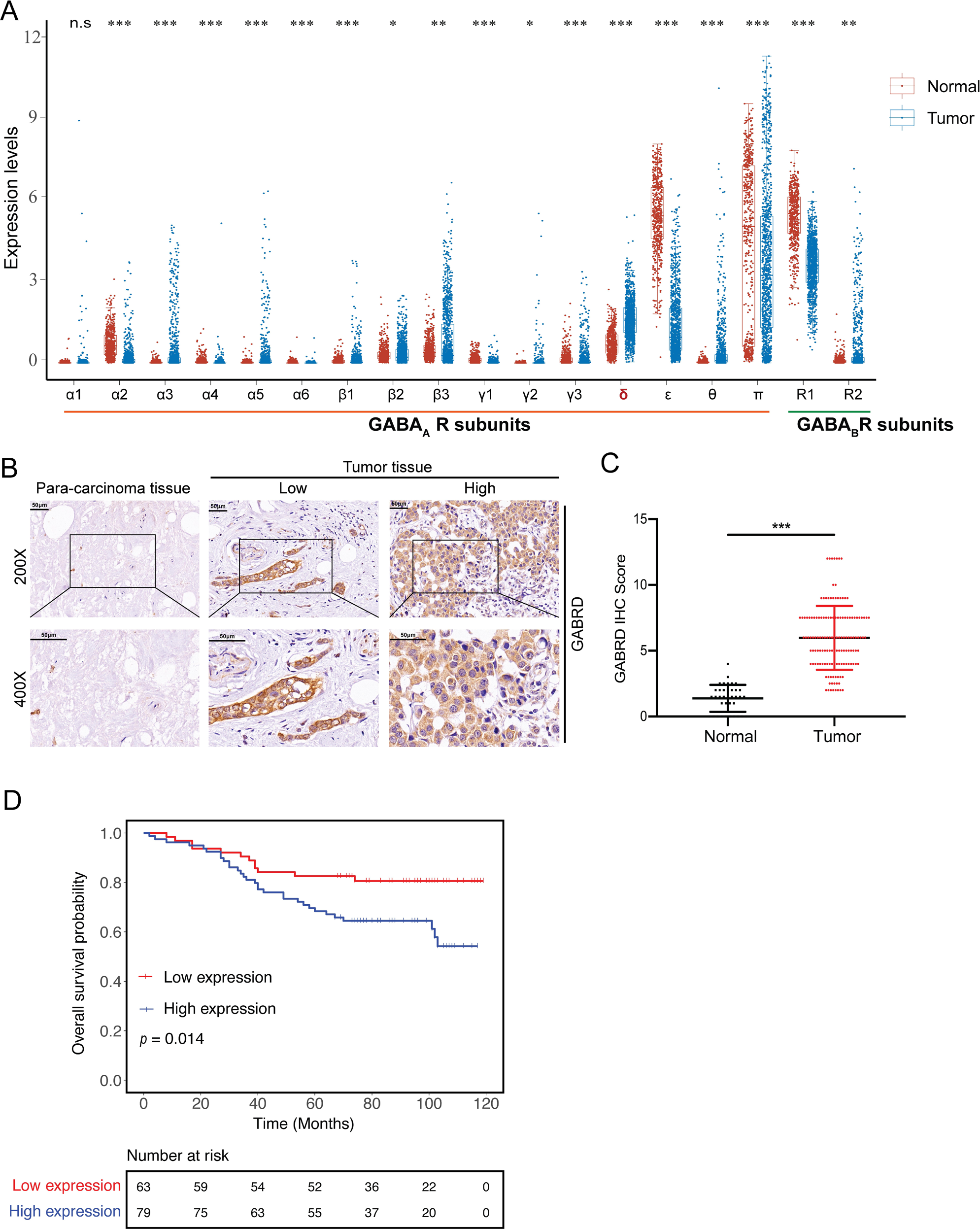
*GABRD* is upregulated in breast cancer and predicts a poor prognosis. **A** Breast cancer expression profile obtained from the TCGA database (blue) and the normal tissue RNA-seq profile obtained from the TCGA database and GTEx database (red). The expression levels of GABA_A_R and GABA_B_R subunits were shown. *GABRD* were most significantly upregulated in breast cancer tissue than normal tissue. \**p<*0.05, \*\**p<*0.01, \*\*\**p<*0.001; **B** Expression of *GABRD* in breast cancer tissues (n=150) and paracarcinoma tissues (n=52) detected by IHC; **C** GABRD IHC score between tumor tissue (tumor) and paracarcinoma tissue (normal). \*\*\**p<*0.001; **D** Kaplan–Meier survival analysis of breast cancer patients with low and high *GABRD* expression levels, p=0.014.

**Table 1.** Expression patterns of GABRD in breast cancer tissues and para-carcinoma tissues revealed in immunohistochemistry analysis, *p*<0.001.

| GABRD expression | Tumor tissue |  | Para-carcinoma tissue |  |
| --- | --- | --- | --- | --- |
|  | Cases | Percentage | Cases | Percentage |
| Low | 63 | 44.4% | 37 | 100% |
| High | 79 | 55.6% | 0 | - |

**Table 2.** Relationship between GABRD expression and tumor characteristics in patients with breast cancer. *p<0.05.

| Features | No. of patients | GABRD expression |  | p value |
| --- | --- | --- | --- | --- |
|  |  | Low | High |  |
| All patients | 142 | 63 | 79 |  |
| Age (years) |  |  |  | 0.096 |
| $\leq 60$ | 70 | 36 | 34 | |
| $> 60$ | 72 | 27 | 45 | |
| Grade |  |  |  | 0.230 |
| I | 1 | 0 | 1 |  |
| II | 70 | 36 | 34 |  |
| III | 61 | 24 | 37 |  |
| Stage |  |  |  | 0.022* |
| 1 | 24 | 16 | 8 |  |
| 2 | 76 | 34 | 42 |  |
| 3 | 37 | 13 | 24 |  |
| T infiltrate |  |  |  | 0.015* |
| T1 | 36 | 23 | 13 |  |
| T2 | 87 | 35 | 52 |  |
| T3 | 13 | 3 | 10 |  |
| T4 | 2 | 2 | 0 |  |
| Lymphatic metastasis |  |  |  | 0.182 |
| N0 | 74 | 36 | 38 |  |
| N1 | 35 | 17 | 18 |  |
| N2 | 19 | 7 | 12 |  |
| N3 | 12 | 3 | 9 |  |
| Tumor size |  |  |  | 0.024* |
| ≤3cm | 58 | 33 | 25 |  |
| >3cm | 80 | 30 | 50 |  |
| Lymph node positive |  |  |  | 0.371 |
| =0 | 70 | 34 | 36 |  |
| >0 | 66 | 27 | 39 |  |

### Knockdown of *GABRD* expression inhibited the proliferation, migration and invasion potential of breast cancer cells and enhanced apoptosis

After cell lines with stable *GABRD* knockdown were constructed using lentiviral-mediated shRNA interference, the effects of *GABRD* knockdown on cell proliferation were assessed by cell counting using Celigo. The numbers of cells in the sh*GABRD* and shCtrl groups within five days after lentivirus infection was recorded over time (Fig. 2A). Compared with the shCtrl group, the sh*GABRD* group showed significantly inhibited cell proliferation in both MDA-MB-231 and BT-549 cell lines (*p<*0.001).

**Figure 2.**
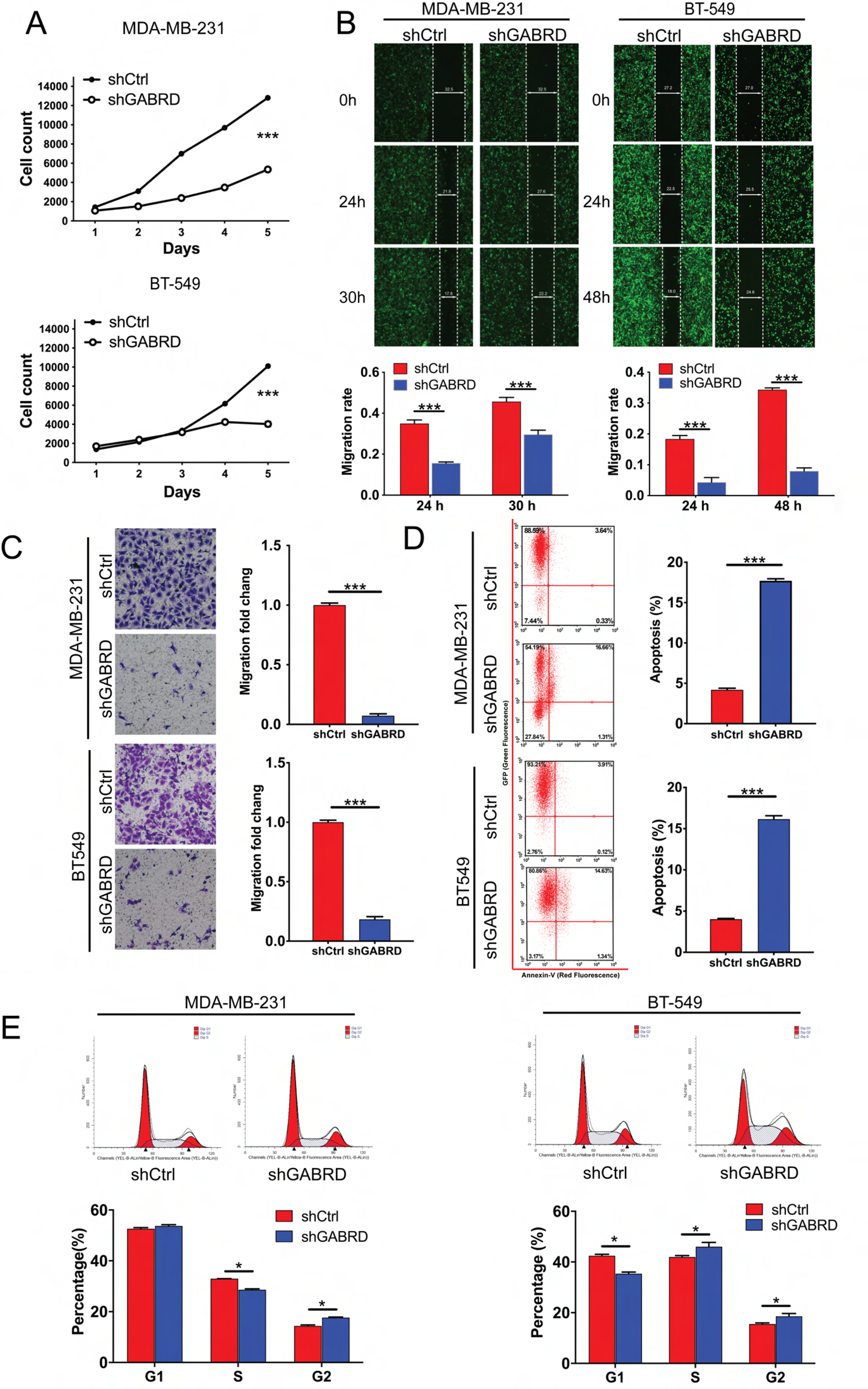
*GABRD* knockdown inhibited breast cancer development *in vitro*. **A** Cell count assay; **B** Wound healing assay; **C** Transwell assay; **D** Rate of apoptosis determined by flow cytometry; **E** Percentage of cells in each phase of the cell cycle determined by flow cytometry. \**p<*0.05, \*\**p<*0.01, \*\*\**p<*0.001.

The wound healing assay and transwell assay were used to measure cell migration and invasion. In the wound healing assay (Fig. 2B), we found that the relative cell migration distance was significantly shorter in the sh*GABRD* group than in the shCtrl group. The 30 h MDA-MB-231 cell migration rate in the sh*GABRD* group was reduced by 35% (*p<*0.001). The 48 h BT-549 cell migration rate in the sh*GABRD* group was reduced by 76% (*p<*0.001). The Transwell metastasis rate in the sh*GABRD* group was decreased by 93% in the MDA-MB-231 cell line (*p<*0.001) and 82% in the BT-549 cell line (*p<*0.001) compared with the shCtrl group (Fig. 2C). This indicated that MDA-MB-231 and BT-549 breast cancer cells exhibited significantly reduced invasive tendencies after *GABRD* knockdown.

Flow cytometry assays were performed to further detect the effects of *GABRD* on apoptosis and the cell cycle distribution. Fig. 2D shows a significant increase in the number of apoptotic cells in the sh*GABRD* group compared with the shCtrl group (*p<*0.001) in both cell lines. Furthermore, the relevant genes in the human apoptosis signaling pathway were also detected using an apoptosis antibody array. Compared with the shCtrl group, the expression of Bax, Caspase3, Fas, p27, p53, and SMAC was significantly higher in the sh*GABRD* group, but the expression of IGF-II, Survivin, sTNF-R1, and XIAP was significantly lower in the sh*GABRD* group (Fig. S1). Cell cycle analysis revealed a significant increase in the number of sh*GABRD* cells in G2 phase compared with the shCtrl group (Fig. 2E). The results indicated that the inhibitory effect of *GABRD* knockdown on cell proliferation was most likely mediated by G2 cell cycle arrest and apoptosis.

### Knockdown of *GABRD* inhibited the tumorigenesis of breast cancer cells *in vivo*

The tumorigenicity of *GABRD* was further tested in a nude mouse assay in which *GABRD* knockdown cells (sh*GABRD*) and normal negative control MDA-MB-231 cells (shCtrl) were inoculated subcutaneously, and tumor growth was assessed. The tumors in the two groups of nude mice are shown in Fig. 3A. The average volume of tumors induced by sh*GABRD* cells was significantly decreased compared with that of shCtrl tumors (*p<*0.001, Fig. 3E). Tumor growth was monitored using luciferase imaging. The quantitative analysis of fluorescence expression showed that expression in the sh*GABRD* group was significantly lower than that in the shCtrl group, and the tumor weights followed the same trend (*p<*0.01, Fig. 3C, D). In addition, H&E staining and Ki-67 staining showed that Ki-67 expression in the tumors of the sh*GABRD* group was lower than that in the shCtrl group (Fig. 3F), which further verified that *GABRD* knockdown inhibited the proliferation of breast cancer *in vivo*.

**Figure 3.**
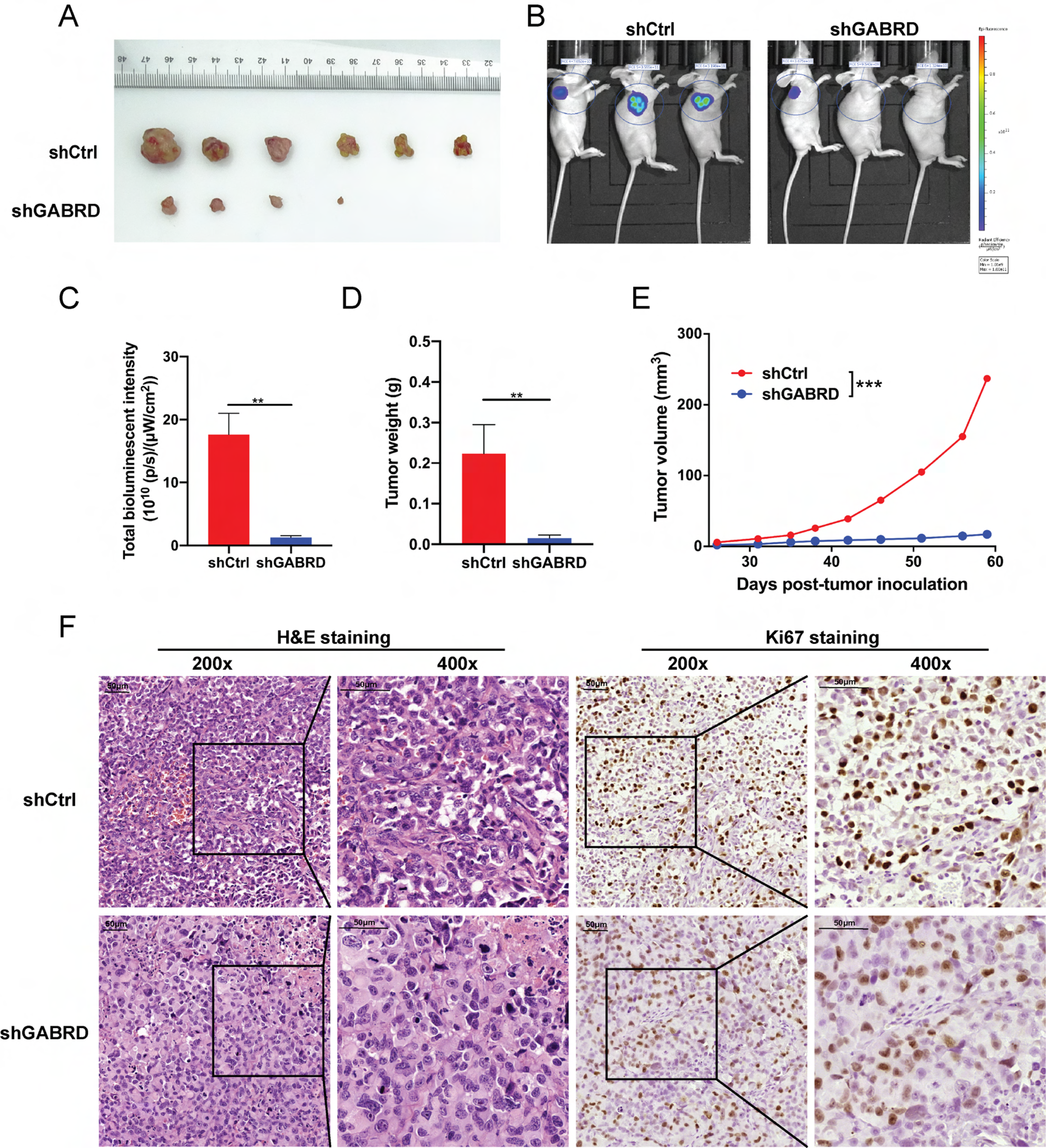
Knockdown of *GABRD* inhibits tumorigenesis and tumor growth of breast cancer *in vivo*. **A** Photographs of tumors excised 59 days after the inoculation of breast cancer cells into nude mice; **B** Fluorescence intensity of tumors detected in animal imaging scanned and **C** quantified as a representation of the tumor burden; **D** Tumor weights were measured after the mice were sacrificed on Day 59; **E** Tumor volume changes with time after inoculation; the data are shown as the mean±SD (n≥3). * p < 0.05, ** p < 0.01, *** p < 0.001.

### GABRD regulates breast cancer progression by modulating CDK1

To further investigate the mechanism by which *GABRD* modulates breast cancer progression, we searched for potential downstream target genes of *GABRD* by IPA analysis based on Affymetrix GeneChip data. In total, 1008 genes were differentially expressed between *GABRD* knockdown MDA-MB-231 cells and normal control cells. After *GABRD* knockdown, 250 genes were upregulated, and 758 genes were downregulated (Fig. 4A). Subsequently, these differentially expressed genes (DEGs) were subjected to enrichment analysis based on the IPA database. Notably, we found that DEGs were more concentrated in cell cycle regulatory signaling pathways, such as G2/M checkpoint regulation and estrogen-mediated S-phase entry (Fig. S2A). The disease and function enrichment of the DEGs also included cancer and the cell cycle (Fig. S2B). This result indicated that the downstream regulation of *GABRD* was closely related to the cell cycle.

**Figure 4.**
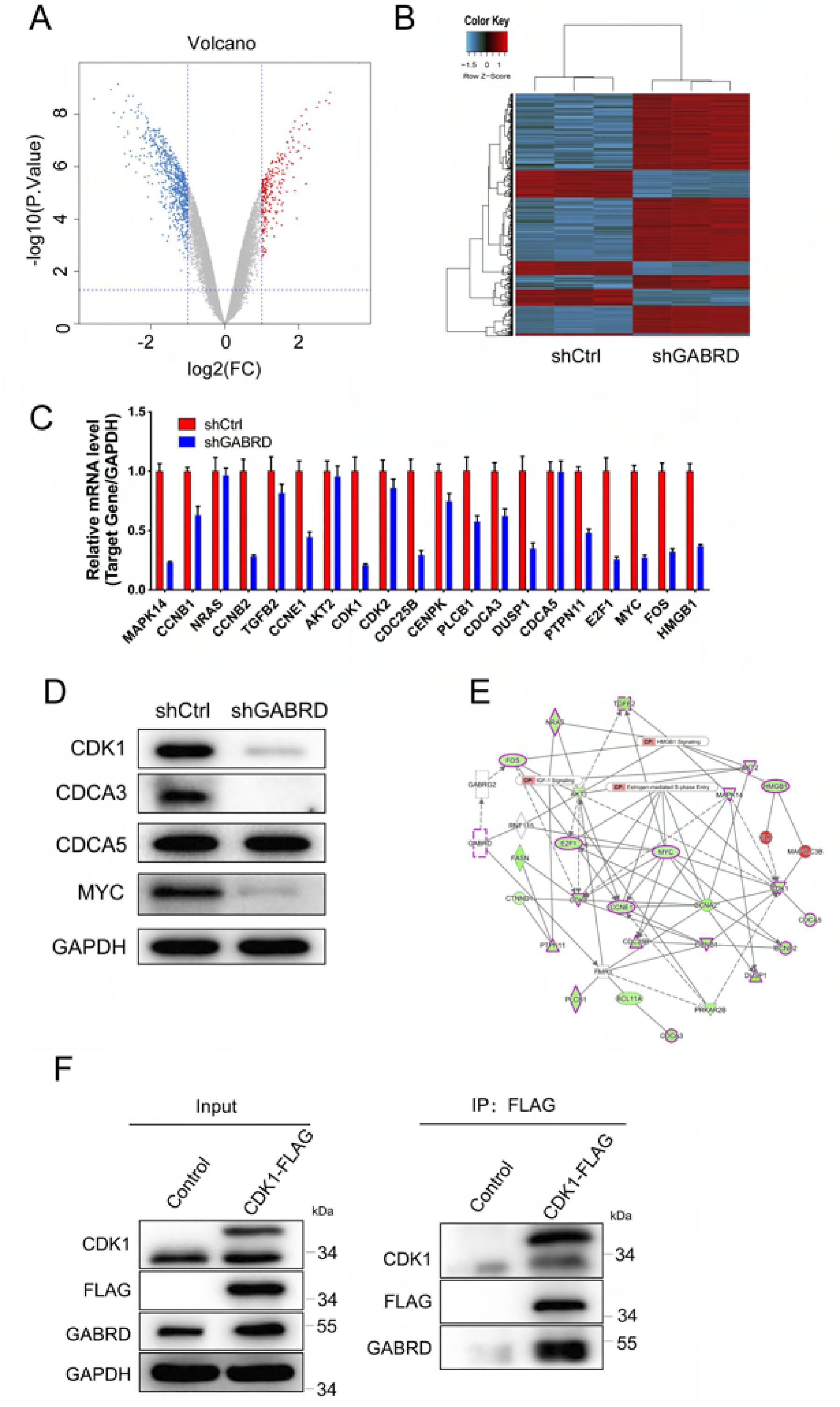
*GABRD* knockdown inhibits breast cancer by interacting with and regulating CDK1. **A** Volcano plot and **B** heatmap showing the differentially expressed genes identified by gene microarray analysis of sh*GABRD* and shCtrl cells. Several differentially expressed genes were selected for further verification by **C** qPCR and **D** western blotting; **E** Molecular interaction network involving *GABRD* and the selected differentially expressed genes constructed based on IPA; **F** Coimmunoprecipitation assay performed to verify the interaction between GABRD and CDK1.

We constructed a network of gene–gene interactions among HMGB1 signaling, estrogen-mediated S-phase entry, the IGF-1 signaling pathway, and other interacting genes with *GABRD* (Fig. 4E). Among these interactions, *GABRD* could indirectly affect HMGB1 signaling, estrogen-mediated S-phase entry, the IGF-1 signaling pathway, and several downstream genes, such as *AKT2* and *CDK1*. Based on the above results, we then concentrated on potential downstream genes associated with *GABRD*-induced breast cancer tumorigenesis using q-PCR (Fig. 4C). Based on the gene expression changes in candidate downstream genes after *GABRD* knockdown, we selected several genes that were significantly decreased, including *CDK1, CDCA3, CDCA5, MYC, ERK, P-ERK, CCND1*, and *PI3KCA*. Next, we used western blotting to separately verify the expression of potential downstream genes. After lentivirus infection, compared with the shCtrl group, the expression of *CDK1, CDCA3*, and *MYC* in the sh*GABRD* group was downregulated, and the expression of *CDCA5* did not change significantly (Fig. 4D). Based on the above results, we assumed that *CDK1* was a potential downstream gene of *GABRD* and conducted a coimmunoprecipitation assay (co-IP) for verification (Fig. 4F). The results showed that the whole-cell lysate contained CDK1 and GABRD proteins. Subsequently, the whole-cell lysates of the control and CDK1-FLAG groups were immunoprecipitated with the FLAG antibody, and the obtained proteins were detected with GABRD, CDK1, and FLAG antibodies. The results showed a clear GABRD band, indicating an interaction between GABRD and CDK1.

### CDK1 plays a crucial role in the GABRD-induced regulation of breast cancer

We constructed a lentiviral expression vector that stably overexpressed *GABRD* in MDA-MB-231 breast cancer cells (*GABRD*+shCtrl), a stable *CDK1* knockdown cell line using lentiviral small hairpin RNAs and the empty vector as a negative control (*shCDK1*+Vector), and a cell line overexpressing GABRD in MDA-MB-231 breast cancer cells with simultaneous *CDK1* knockdown (GABRD+*shCDK1*). We used the empty vector-expressing lentivirus as a control (shCtrl+Vector).

From Fig. 5A, we found that compared with the shCtrl+Vector group, the cells in the *GABRD*+shCtrl group exhibited a faster proliferation rate (*p<*0.001), and *CDK1*-knockdown cells exhibited a slower proliferation rate (*p<*0.01) and could reverse the proliferation effect of *GABRD*. The cell migration and invasion abilities were detected by wound healing tests (Fig. 5C) and transwell tests (Fig. 5D). The degree of cell apoptosis was analyzed by flow cytometry (Fig. 5B). We found that *CDK1* knockdown could partially reverse the promoting effect of *GABRD* on the development of breast cancer.

**Figure 5.**
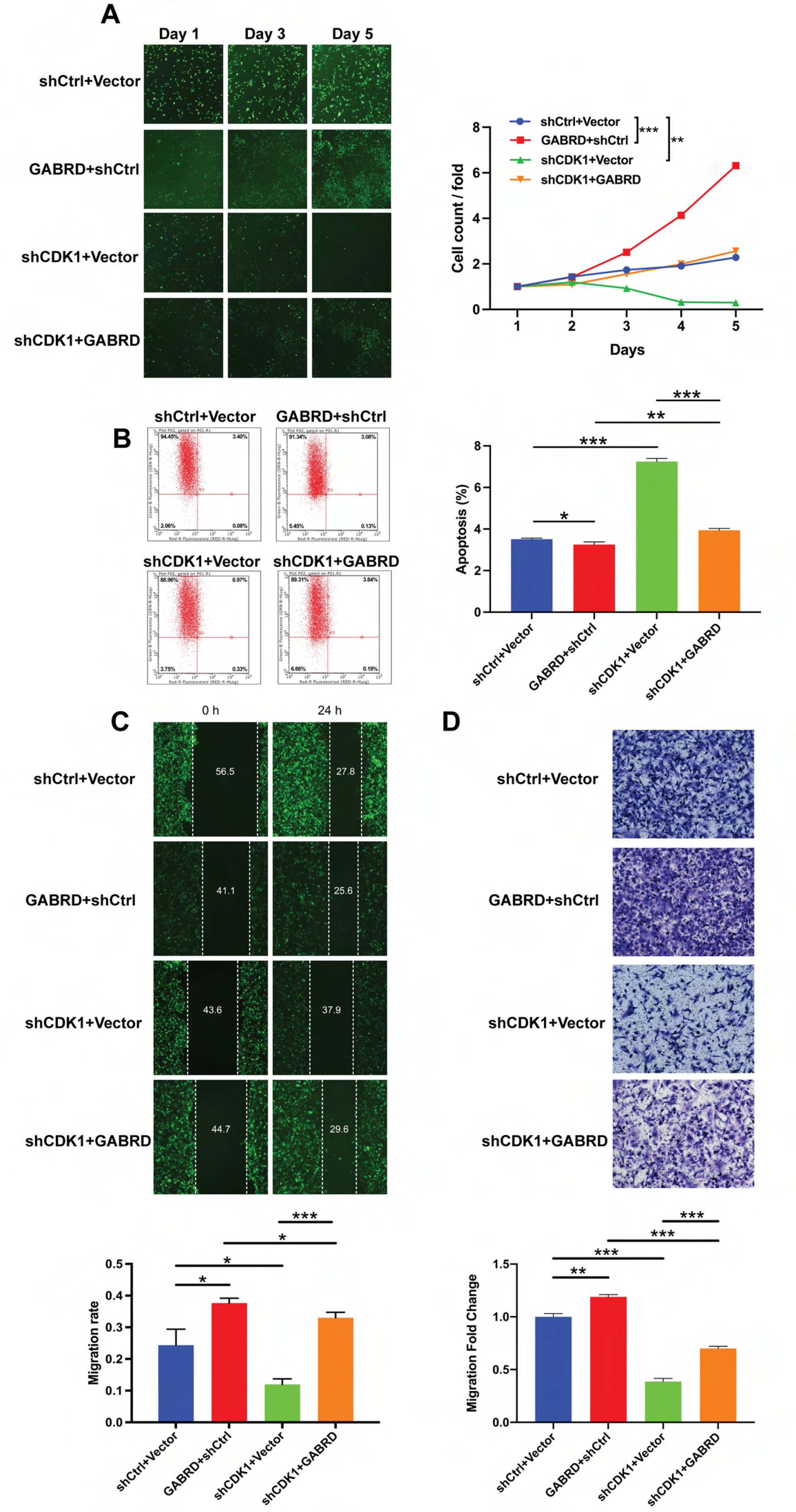
*GABRD*-regulated tumor inhibition is CDK1 dependent. **A** Cell counting assay; **B** Apoptosis analyzed by flow cytometry; **C** Wound healing assay; **D** Transwell assay; the data are shown as the mean±SD (n≥3). * *p<*0.05, ** *p<*0.01, *** *p<*0.001.

Furthermore, we performed a rescue experiment for *in vivo* validation in a nude mouse orthotopic breast tumor model (Fig. 6). We constructed *GABRD* knockdown (sh*GABRD*+Vector), *CDK1* overexpression (*CDK1*+shCtrl), rescue (sh*GABRD*+*CDK1*) and control (shCtrl+Vector) MDA-MB-231 cell lines. Six weeks after the injection, the tumor weights and growth curves of the tumor volume were measured (Fig. 6B, C). The results not only indicated that *CDK1* overexpression could promote tumorigenesis progression but also confirmed that *CDK1* overexpression rescued *GABRD* knockdown.

**Figure 6.**
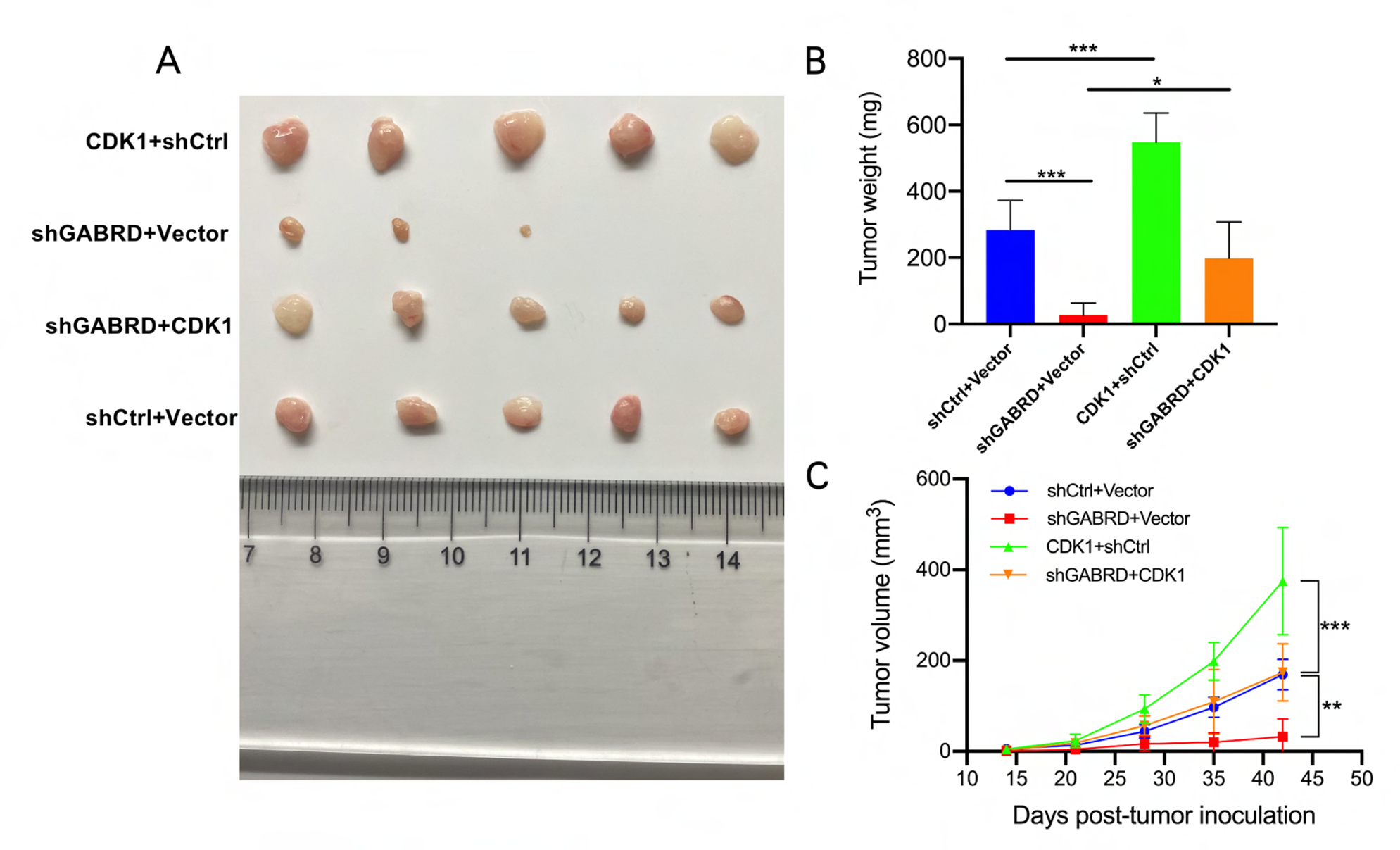
*CDK1* overexpression reverses the inhibition of breast cancer induced by *GABRD* knockdown in an orthotopic mouse model. **A** Photographs of tumors excised 42 days after the inoculation of breast cancer cells into nude mice; **B** Tumor weight measured after sacrifice; **C** Tumor volume changed with time after inoculation; the data are shown as the mean±SD (*n*≥3). * p < 0.05, ** p < 0.01, *** p < 0.001.

### GABRD modulates CDK1 to promote cell proliferation by influencing the cell cycle and G2/M transition

As mentioned above, *GABRD* showed a great impact on the cell cycle, and we performed cell cycle analysis to further verify the mechanism by which the GABRD–CDK1 axis affects breast cancer progression. Flow cytometry was used to determine the cell counts in each phase. We found that compared with the shCtrl+Vector group, the cells in G2 phase in the *GABRD*+shCtrl group were decreased markedly (*p<*0.05), while more cells were arrested in G2 phase in the *shCDK1*+Vector group (*p<*0.01). A moderate change in the *GABRD*+*shCDK1* group was found (Fig. S3B).

## Discussion

The GABA_A_ receptor is a ligand-gated ion channel receptor that mediates rapid synaptic transmission inhibitory effects under physiological conditions. Various neuropsychiatric diseases are associated with GABA signaling, such as schizophrenia, depression, epilepsy, and posttraumatic stress disorder syndrome. GABRD is the δ subunit of the GABA_A_ receptor, and its main function is still under investigation. Most studies have shown that *GABRD* is closely related to the occurrence of various neurological and mental disorder-related symptoms and diseases(5). Recently, some studies have reported the *GABRD* gene expression levels are involved in the development of several malignancies. In particular, the mutation and overexpression of *GABRD* can cause the proliferation of tumor cells, especially in kidney renal clear cell carcinoma, thyroid carcinoma, and breast invasive carcinoma(9). Andrew M. Gross reported that increased *GABRD* levels that could lead to functional changes in the GABA_A_ receptor might play a role in tumor cell differentiation(10). Despite these studies, the role and mechanism of *GABRD* in breast cancer are still unknown. Our study, for the first time, revealed the relationship between *GABRD* and breast cancer.

In this study, we gained insight into the difference in *GABRD* gene expression between tumor tissues and normal tissues in breast cancer patients and its correlations with clinicopathological data (Fig. 1, Table 1). The tissue microarray results showed that *GABRD* was overexpressed in breast cancer tissues, which was also closely related to clinical progression and survival. To further verify our conclusion, we analyzed breast cancer and paracarcinoma tissue expression data from the TCGA database and compared the expression levels of *GABRD*, and the results also indicated that *GABRD* was upregulated in breast cancer tissues (Fig. 1C). Overexpression of *GABRD* might be related to aggressive tumor features, such as tumor size and clinical stage (Table 2). Therefore, it significantly affects the prognoses and survival of patients (Fig. 1D)

The effects of *GABRD* on the proliferation, migration, and invasion of breast cancer were explored (Fig. 2, 3). After *GABRD* knockdown, breast cancer cells exhibited suppressed proliferation and increased apoptosis. The expression of apoptosis-related molecules, such as Bax, p27, and p53, also showed significant differences. In addition, *GABRD* knockdown inhibited the migration and invasion of breast cancer cells, suggesting a role in breast cancer metastasis (Fig. 2). The critical role of *GABRD* was also confirmed in a xenograft mouse model test, which showed a decreased tumor burden according to tumor size and fluorescence analyses in the *GABRD*-knockdown group (Fig. 3). In summary, we demonstrated that *GABRD* expression was associated with malignant behaviors of breast cancer cells. *GABRD* could contribute to the development and progression of breast cancer through various factors.

Subsequently, we explored the mechanisms and key downstream genes potentially regulated by *GABRD*. Through Affymetrix GeneChip analysis and IPA analysis, several potential downstream genes of *GABRD* were selected and then verified by q-PCR and western blotting (Fig. 4). We found that *CDK1* was tightly associated with *GABRD* expression and might act as a key downstream gene of *GABRD* to regulate breast cancer cell processes. Thereafter, the co-IP test also showed a direct interaction between CDK1 and GABRD. These results revealed that CDK1 acts as a downstream molecule of GABRD and exerts its tumor-regulating function.

The *GABRD*-induced enhancement of proliferation and migration was abrogated by the knockdown of *CDK1*, indicating that *GABRD* regulated tumor characteristics in breast cancer by affecting *CDK1*. For further verification, a rescue experiment using a nude mouse orthotopic breast tumor model was performed. The tumor-suppressing effect of *GABRD* knockdown was rescued by *CDK1* overexpression. The *in vivo* experiment strongly suggested that the interaction of *GABRD* with *CDK1* affected breast cancer development.

CDK1 is a highly conserved protein that functions as a serine/threonine kinase. Through the phosphorylation of a diverse set of substrates, including apoptosis and cell cycle regulators, by CDK1 complexes and their cyclin partners, CDK1 can regulate transitions through distinct cell cycle phases(11). Previous studies have indicated that *CDK1* overexpression promotes the oncogenesis and progression of several cancers, and *CDK1* inhibition could lead to cell cycle arrest and apoptosis(12, 13). In addition, Kim et al. found that breast cancer patients with high CDK1 and CDK2 activity had markedly reduced 5-year relapse-free survival compared to breast cancer patients with low CDK1 and CDK2 activity(14). Moreover, as proposed by Liu et al., CDK1 inhibition has a therapeutic effect on triple-negative breast cancer by inhibiting tumor growth and inducing cell apoptosis(15), implying that *CDK1* can serve as a potential key gene affecting the outcomes of breast cancer patients. The CDK1/cyclin B1 complex is crucial in regulating the G2-M phase transition based on previous studies(16). Thus, we assumed that *GABRD* could promote tumor proliferation by regulating CDK1 and accelerating the G2/M phase transition. This was verified by flow cytometry detection of the cell cycle (Fig. 2E, S3). More cells were arrested in G2 phase after *GABRD* knockdown. In addition, the number of cells in G2 phase was significantly reduced after *GABRD* overexpression, which could be restored by *CDK1* knockdown. In summary, *GABRD* promoted breast cancer proliferation by directly regulating CDK1 to affect the cell cycle, especially in the G2/M phase transition.

Currently, there are many studies and applications of CDK4/6 inhibitors in breast cancer treatment, which can effectively block progression of tumor cells from G1 phase to S phase(17, 18). CDK4/6 inhibitors, such as Palbociclib, are already available on the market. There are few studies on CDK1 inhibitors for treating breast cancer, leaving a broad gap. CDK1 inhibitors can play a significant role in specific tumor subtypes. For instance, MYC-dependent breast cancer cells were sensitive to CDK1 inhibition, excluding CDK4/6 or CDK2(19), and aggressive breast tumors with c-myc overexpression were more sensitive to CDK1/2 inhibitors than CDK4/6(20). Therefore, CDK1 inhibitors are worthy of further study as potential therapies for specific breast cancer subtypes. As an effective target, GABRD synergistic therapy is expected to promote CDK1 inhibitor function.

In conclusion, our research provides new insights into understanding *GABRD*-mediated CDK1 activation and promotion in breast cancer tumorigenesis. *GABRD* can provide a target for further cell cycle-related therapy development and a potential enhancement of the anticancer effects of CDK1 inhibitors. Further studies regarding the role of *GABRD* and its specific downstream pathway based on CDK1 would be worthwhile. Bitarget combination therapy based on GABRD and CDK1 could also prove helpful based on possible animal and clinical research in the future.

## Material and Methods

### Ethical approval

Acquisition and experiments on human samples were performed according to the Helsinki Declaration and ethical approval was obtained from the Ethics Committee of National Cancer Center (NCC) (Approval number: 20/468-2664). Animal ethics approval was granted by Animal Ethics Committee of NCC (Approval number: NCC2021A220).

### Patient samples

In total, 150 breast cancer tissue and 52 paracarcinoma tissue samples were obtained from the Department of Pathology in NCC. These samples were used to determine *GABRD* expression levels through tissue microarrays. Clinical information about the patients was obtained from electronic medical records, while follow-ups were conducted through phone calls or electronic medical record searches.

### Cell culture and transfection

The human breast cancer cell lines MDA-MB-231 and BT-549 were obtained from the Cell Bank of Sciences (Shanghai, China). The cell culture conditions were DMEM+10% FBS and 100 μg/ml penicillin–streptomycin at 37 °C and 5% CO2. Stably transfected cell lines were created by constructing *GABRD* or *CDK1* short-hairpin RNA lentiviral vectors (BR-V108) and *GABRD*- or *CDK1*-overexpression lentivirus vectors (LV-007). The sequences used are listed in Table S2. Puromycin was used to screen for positive colonies over 4 weeks.

### Tumor-forming experiment (Xenograft models)

BALB/c nude mice (4 weeks old, 18±2.5 g) were purchased from Beijing Vital River Laboratory Animal Technology Co., Ltd. The experimental animal protocol was approved by the Ethics Committee of NCC. For the functional verification experiments, two groups were established (shCtrl group and sh*GABRD* group), and 6 mice in each group received 4 x 10^6^ sh*GABRD*- or shCtrl-transfected MDA-MB-231 cells with Matrigel by subcutaneous injection. The tumor length and width were measured every week using caliper. After 60 days, a Xenogen IVIS imaging system (IVIS Spectrum, Perkin Elmer) was used for live animal imaging. Then, the mice were sacrificed by carbon dioxide asphyxiation, and the tumors were harvested. In the gene rescue experiments, four groups were established (shCtrl+Vector, sh*GABRD*+Vector, *CDK1*+shCtrl, and sh*GABRD*+*CDK1*). A total of 4×10^6^ MDA-MB-231 cells transfected with Matrigel were orthotopically injected into the first mammary fat pod of the five mice in each group. The tumor length and width were measured every week using calipers. After 42 days, the mice were sacrificed by carbon dioxide asphyxiation for tumor excision and measurement.

### Tissue microarray

IHC analysis of GABRD protein expression was performed on tissue microarrays produced from 150 breast cancer tissues and 52 paracarcinoma tissues. The relevant antibody information is shown in Table S1. The IHC results were scored using a modified H-score system, which combined the staining intensity score and the staining percentage score.

### Cell counting assay

Cells in the logarithmic phase were seeded into 96-well plates (2000 cells per well). The plates were continuously quantified using Celigo (Nexcelom) over the following five days.

### Cell apoptosis assay

Cell apoptosis was analyzed using flow cytometry (Millipore, USA). An Annexin V-APC apoptosis detection kit (eBioscience, San Diego, CA, USA) was used. Cells were harvested to produce cell suspensions containing 1 × 10^6^–1 × 10^7^/mL cells using binding buffer. Then, 5 μL annexin V-APC was added to 100 μL cell suspensions and incubated at 25 °C for 10–15 min. Then, FACS was used for cell apoptotic analyses.

### Wound healing assay

MDA-MB-231 or BT-549 cells were seeded into a 6-well plate after being transfected with lentiviruses. A single layer of cells was scratched in a straight line using a 10 µl sterile pipette tip and washed with PBS. The migration area was analyzed by Cellomics as follows: migration area = X h cell area −0 h cell area.

### Transwell assay

A transwell kit (3422 Corning) was used to perform the transwell assay. Then, 100 µL MDA-MB-231 or BT-549 cell suspension in 30% fetal bovine serum (FBS) medium was added to the upper chamber, and the same was performed for the lower wells. The plate was placed in a tissue culture incubator (37 ℃, 24 h). The medium was wipd off of the nontransferred cells in the upper chamber. Subsequently, cells that migrated to the lower surface of the membrane were stained with crystal violet and counted under a microscope after fixation with 4% paraformaldehyde.

### Western Blot

The total cellular proteins were obtained from the above cells using a Total Protein Extraction Kit (KeyGEN, China), and the protein concentrations were quantified using the BCA method. Then, related expressed molecules were determined by western blotting. Sodium dodecyl sulfate–polyacrylamide gel electrophoresis (SDS–PAGE) was used to separate the target proteins. The proteins were transferred to polyvinylidene fluoride (PVDF) membranes. Five percent nonfat milk was used to block the nonspecific sites. Then, the membranes were incubated with the primary antibody overnight at 4 °C, followed by incubation with the secondary antibody. The antibodies used are shown in Table S1. After adding the luminescent reagent, the film was exposed.

### Real-time qPCR

Total RNA was first extracted using the TRIzol method. One microgram of RNA was reverse-transcribed into cDNA using the Hiscript QRT Supermix for qPCR synthesis kit (Vazyme). All DNA and RNA concentrations were measured by a NanoDrop 2000 spectrophotometer (Thermo). qPCR was performed using SYBR Green Mastermixes (Vazyme) in a 10-µl reaction system. The primers used are listed in Table S3. Amplification was performed in 2 steps of 45 cycles. Final fluorescence data were analyzed using the 2^(-ΔΔCt)^ method (21).

### Cell cycle assay

The cell cycle was analyzed using flow cytometry (Millipore, USA) with the Cell Cycle Assay Kit (Beyotime). Cells were digested and collected. Then, 70% ethanol was used to fix the cells overnight. Afterward, the cells were washed twice with PBS and suspended in propidium iodide (PI) buffer before cell cycle detection.

### Co-IP

For co-IP, MDA-MB-231 cells were lysed with RIPA lysis buffer on ice. The BCA method was used to determine the protein amounts. Cell lysates were collected and incubated with the antibodies overnight, followed by an additional incubation with protein A/G magnetic beads for 2 h. The antibodies used are shown in Table S1. The complexes were washed with washing buffer and eluted with elution buffer for western blotting.

### Human apoptosis antibody array

A Human Apoptosis Antibody Array kit was purchased from Abcam (ab134001), and this assay can simultaneously detect 43 human apoptosis-related proteins. Total proteins were extracted after adding cell lysis buffer. The protein amounts were quantified using the BCA method. The total proteins were incubated with biotin-conjugated anti-cytokines overnight at 4°C. After washing, streptavidin-HRP incubation was performed. Enhanced chemiluminescence (ECL) was used to visualize protein spots.

### Microarray

Microarray analysis was performed using the Affymetrix GeneChip Primeview Human Gene Expression Array. Total RNA was extracted from sh*GABRD*- or shCtrl-transfected MDA-MB-231 cells using the TRIzol method. cRNA was prepared from total RNA and hybridized onto a Gene Expression Array. Subsequent data processing was performed with *R software* (R verson 3.5.2) and *Ingenuity Pathway Analysis software* (IPA, Ingenuity Systems) for differentially expressed gene analysis, classical signal pathway analysis, and disease and functional analyses.

### Statistical analyses

Statistical analysis was performed using SPSS software (SPSS 17.0). Statistical analysis was performed using Student’s t test for continuous variables. Categorical data were tested using the χ^2^ test or the continuity correction χ^2^ test. Survival curves were plotted using the Kaplan–Meier method (log-rank test). A Bonferroni post hoc test was used for pairwise comparisons. *p<*0.05 indicated statistically significant differences.

## Statements and Declarations

### Ethical Approval and Consent to participate

Acquisition and experiments on human samples were performed according to the Helsinki Declaration and ethical approval was obtained from the Ethics Committee of National Cancer Center (NCC, 20/468-2664). Informed consents were collected. Experiments on mice in this study were performed in accordance with institutional, national, or international guidelines. The experimental animal protocol was approved by the Ethics Committee of NCC (NCC2021A220).

### Availability of supporting data

All data generated or analyzed during this study are included in this published article and its supplementary information files.

### Authors’ contributions

X.W. conceived, designed, coordinated and directed this study. Q.S., R.F., K.F performed all molecular and animal experiments. C.Y., S.Z., J.L. supervised and performed the statistical analysis in this study. Q.S. wrote this manuscript. X.K., J.Y., R.Z., X.M., Xiang W. assessed and modified the manuscript. All authors reviewed and approved the final manuscript.

## Acknowledgements

Not applicable

